# Adaptive immune determinants of viral clearance and protection in mouse models of SARS-CoV-2

**DOI:** 10.1101/2021.05.19.444825

**Authors:** Benjamin Israelow, Tianyang Mao, Jonathan Klein, Eric Song, Bridget Menasche, Saad B. Omer, Akiko Iwasaki

**Affiliations:** Department of Immunobiology, Yale University School of Medicine, New Haven, CT, USA; Department of Medicine, Section of Infectious Diseases, Yale University School of Medicine, New Haven, CT, USA; Department of Laboratory Medicine, Yale University School of Medicine, New Haven, CT, USA; Department of Epidemiology of Microbial Diseases, Yale School of Public Health, New Haven, CT, USA; Yale Institute for Global Health, Yale University, New Haven, CT, USA; Howard Hughes Medical Institute, Chevy Chase, MD, USA

## Abstract

Severe acute respiratory syndrome coronavirus 2 (SARS-CoV-2) has caused more than 160 million infections and more than 3 million deaths worldwide. While effective vaccines are currently being deployed, the adaptive immune determinants which promote viral clearance and confer protection remain poorly defined. Using mouse models of SARS-CoV-2, we demonstrate that both humoral and cellular adaptive immunity contributes to viral clearance in the setting of primary infection. Furthermore, we find that either convalescent mice, or mice that receive mRNA vaccination are protected from both homologous infection and infection with a variant of concern, B.1.351. Additionally, we find this protection to be largely mediated by antibody response and not cellular immunity. These results highlight the *in vivo* protective capacity of antibodies generated to both vaccine and natural infection.

**One-Sentence Summary:** Defining the roles of humoral and cellular adaptive immunity in viral clearance and protection from SARS-CoV-2 and a variant of concern.

## Main Text

The development of adaptive immune responses to natural SARS-CoV-2 infection as well as to SARS-CoV-2 vaccination in people have been well characterized (*1–4*). Additional studies have reported that vaccine-induced, and natural-infection induced immunity in animal models (non-human primates (*5–9*), mice (*10, 11*), and hamsters (*12*)) are sufficient for protection from homologous SARS-CoV-2 challenge. Phase III vaccine clinical trials as well as post-marketing vaccine effectiveness studies provide evidence of the development of protective immunity in humans (*13, 14*). Epidemiologic studies of natural infection have also indicated that adaptive immune memory is sufficient to protect against SARS-CoV-2 reinfection in the majority of cases (*15*). However, few of these studies have identified the essential correlates of protective adaptive immunity. The relative contributions of cellular and humoral immunity in the clearance of SARS-CoV-2 and protection from reinfection remain poorly defined in both vaccine-induced immunity as well as natural infection-induced immunity. Importantly, questions regarding the degree to which T cell mediated immune memory may compensate for waning antibody mediated immunity have become even more critical given the emergence of viral variants of concern (VOC) that evade neutralizing antibody responses from both vaccinated and convalescent individuals (*16–19*).

To address these questions surrounding the immunological determinants of protection from SARS-CoV-2 infection, we utilized our previously reported mouse model which employs *in vivo* transduction of the mouse respiratory tract with adeno-associated virus expressing human ACE2 (AAV-hACE2) (Fig 1a) (*20*). This model enables us to infect mice of varying genetic backgrounds to identify how deficiencies in specific components of the adaptive immune system affect clearance of and protection from SARS-CoV-2. First, to assess whether adaptive immunity is required for clearance of SARS-CoV-2 upon primary infection, we administered AAV-hACE2 to Rag1^-/-^ mice, which do not produce any mature B or T lymphocytes. These mice were infected with 1×10^6^ PFU of SARS-CoV-2 WA1 strain (USA-WA1/2020). Interestingly, we found that these mice became persistently infected and were unable to clear the virus, maintaining stable levels of viral RNA and infectious virus for greater than 14 days post-infection in lung tissues (DPI) (Fig 1B,C). This is in contrast to C57Bl/6J (WT) mice, which cleared the virus by 7 DPI as indicated by loss of culturable SARS-CoV-2 by plaque assay and reduction of viral RNA by RT-PCR (Fig 1B,C). While our previous work suggested that innate immunity played a minimal role in clearing SARS-CoV-2 during acute infection in mice (*20*), these data confirm that in the absence of adaptive immunity, the innate immune responses are insufficient to clear acute SARS-CoV-2 infection. These results are consistent with case reports describing patients with either genetic or acquired defects in their adaptive immunity, who are unable to clear SARS-CoV-2 (*21–23*). Next, to assess the requirement of humoral immunity in SARS-CoV-2 clearance, we infected AAV-hACE2 μMT mice (mice deficient in B lymphocytes), and found that these mice retained the ability to clear SARS-CoV-2. However, it occurred more slowly than in WT infected mice, as indicated by the presence of infectious virus at 7 DPI (4/8 vs 0/4) and 17x more viral RNA at 14 DPI (Fig 1B,C). These results suggest that cellular immunity is sufficient for viral clearance in the setting of acute infection, even in the absence of humoral responses; however, humoral immunity is not completely redundant in its contribution to viral clearance.

**Figure 1:**
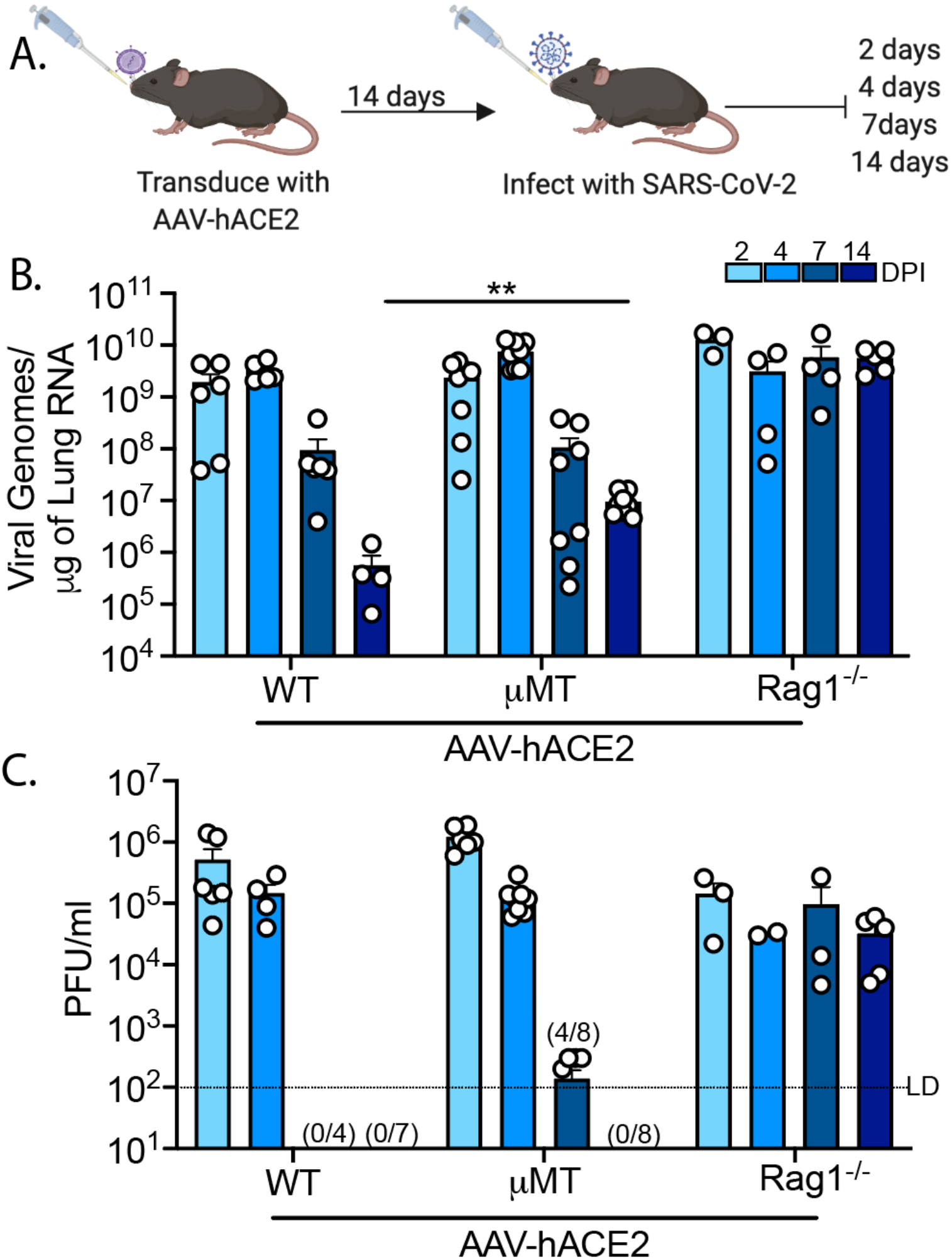
SARS-CoV-2 clearance in RAG^-/-^ and mMT mice. (A) Experimental schematic. C57Bl/6J (WT), B6.129S7-Rag1tm1Mom/J (Rag^-/-^), B6.129S2-Ighmtm1Cgn/J (μMT) mice are intratracheally transduced with 10^11^ GC (genomic copies) of AAV encoding human ACE2 (AAV-hACE2) and allowed to recover for 14 days. Mice are then infected with WA1 strain SARS-CoV-2 at 10^6^ PFU intranasally and lungs samples are collected at 2, 4, 7, and 14 day post infection and assessed by (B) quantitative PCR and (C) plaque assay. Values noted on X-axis (C) indicate numbers of samples tested positive/number of samples. LD (limit of detection). Individual values noted as dots and bars as mean ± SEM from 3-8 samples of two-three independent experiments. P values were calculated by two-tailed Mann-Whitney test. *, P< 0.05; **, P < 0.01; ***, P < 0.001, ****, P < 0.001

To further delineate the role of T cell subsets required for viral clearance, we infected AAV-hACE2 WT mice that were treated with αCD4, αCD8, or αCD4+αCD8 depleting antibodies (Fig 2A). We found that depletion CD8^+^ T cells alone each inhibited viral clearance, as indicated by 7.4x more RNA than PBS control at 14DPI (Fig 2B). We also found that depletion of CD4^+^ T cells alone inhibited viral clearance by 6.2x compared to control at 14 DPI (Fig 2B). Depletion of both CD4^+^ and CD8^+^ T cells had a synergistic effect leading to a 31x increase in viral RNA persistence at 14 DPI (Fig 1B). These levels were similar to that seen in AAV-hACE2 Rag^-/-^ infected mice (Fig 1B). Notably, CD4^+^ T cell depletion significantly reduced the development of anti-SARS-CoV-2 Spike S1 and RBD antibody titers, consistent with CD4 helper function in antibody development against SARS-CoV-2 Spike and RBD (Fig 2D,E). These results also indicate that while adaptive immunity is required, either humoral or cellular adaptive immunity on its own is sufficient for viral clearance during a primary infection.

**Figure 2:**
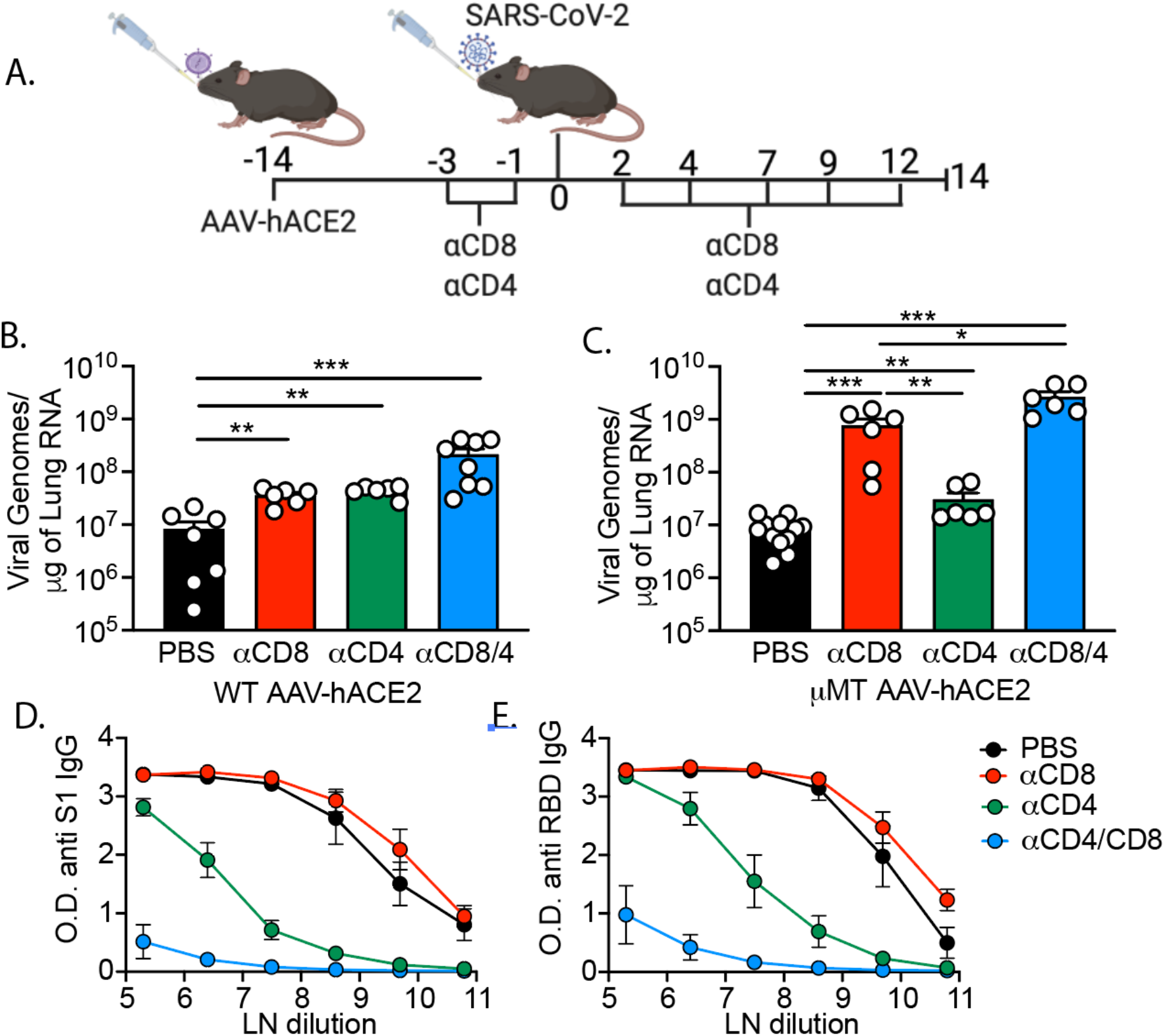
SARS-CoV-2 clearance in CD4^+^ and CD8^+^ T cell depleted WT and mMT mice. (A) Experimental design. AAV-hACE2 transduced WT (B) and mMT (C) were injected (intraperitoneal) with 200μg anti-mouse CD8a (clone 2.43) or 200μg anti-mouse CD4 (clone GK1.5) starting at 3 days prior to infection and given every 2 to 3 days as indicated until 12 DPI. Mice were infected with WA1 strain SARS-CoV-2 at 10^6^ PFU intranasally and lungs were collected at 14 DPI and assessed by quantitative PCR. Individual values noted as dots and bars as mean ± SEM from 6-12 samples of two-three independent experiments. Anti-SARS-CoV-2 antibody concentration measured from serum dilutions from (B) against (D) Spike S1 and (E) receptor binding domain (RBD), values natural log (LN) transformed noted as mean ± SEM from 3-4 samples of 2 independent experiments. P values were calculated by two-tailed Mann-Whitney test. *, P< 0.05; **, P < 0.01; ***, P < 0.001, ****, P < 0.001

Next, we infected AAV-hACE2 μMT mice that had been treated with αCD4, αCD8 or αCD4+αCD8 depleting antibodies to assess the T cell component required for viral clearance in the absence of humoral responses. We found that AAV-hACE2 μMT mice treated with CD8-depleting antibody maintained significantly higher levels of viral RNA (92x) than the PBS controls (Fig 2C). On the other hand, AAV-hACE2 μMT mice treated with αCD4 antibody alone sustained only a small yet significant (3.7x) increase in viral RNA relative to control at 14DPI (Fig 2C). Depletion of both CD4 and CD8 T cells lead to significantly higher levels of viral RNA (320x) compared to PBS treared at 14 DPI (Fig 2C). Interestingly, similar to the difference between αCD4 treated, and PBS treated mice, there was a small increase in viral RNA in αCD4+αCD8 treated μMT mice relative to αCD8 treated mice (3.5x), suggesting a small role of CD4^+^ T cells in the cellular immunity component of viral clearance in the absence of its effect on antibody development. Taken together, these data indicate that CD8 T cells are sufficient for viral clearance in the absence of humoral immunity, and that the major function of CD4 T cells in clearance of acute SARS-CoV-2 is in instructing humoral immunity rather than assisting cellular immunity during primary infection.

Next, we investigated the sufficiency of SARS-CoV-2 specific T cell mediated vs. antibody mediated immunity to protect against SARS-CoV-2 infection. To this end, we performed adoptive transfer of either T cells or sera from previously infected (convalescent) mice (Fig 3A). We infected AAV-hACE2 WT mice with 1×10^6^ PFU SARS-CoV-2. At 14 DPI, sera was harvested and total T-cells from mediastinal (draining) lymph nodes were isolated via magnetic negative selection. Recipient AAV-hACE2 Rag1^-/-^ mice were intravenously injected with either 2×10^6^ T cells or with 200μl of pooled serum. At 24 hours post-transfer, mice were infected with 1×10^6^ PFU SARS-CoV-2 intranasally. At 7 DPI, we found that while convalescent T cell transfer resulted in detectable reduction in viral RNA and titers in the lung, transfer of convalescent sera resulted in complete reduction in infectious viral load in the lung (Fig. 3B, C). These results indicated that in isolation, either cellular or humoral immunity can significantly reduce homologous SARS-CoV-2 infection, with antibodies being able to completely eliminate infectious virus from the lung.

**Figure 3:**
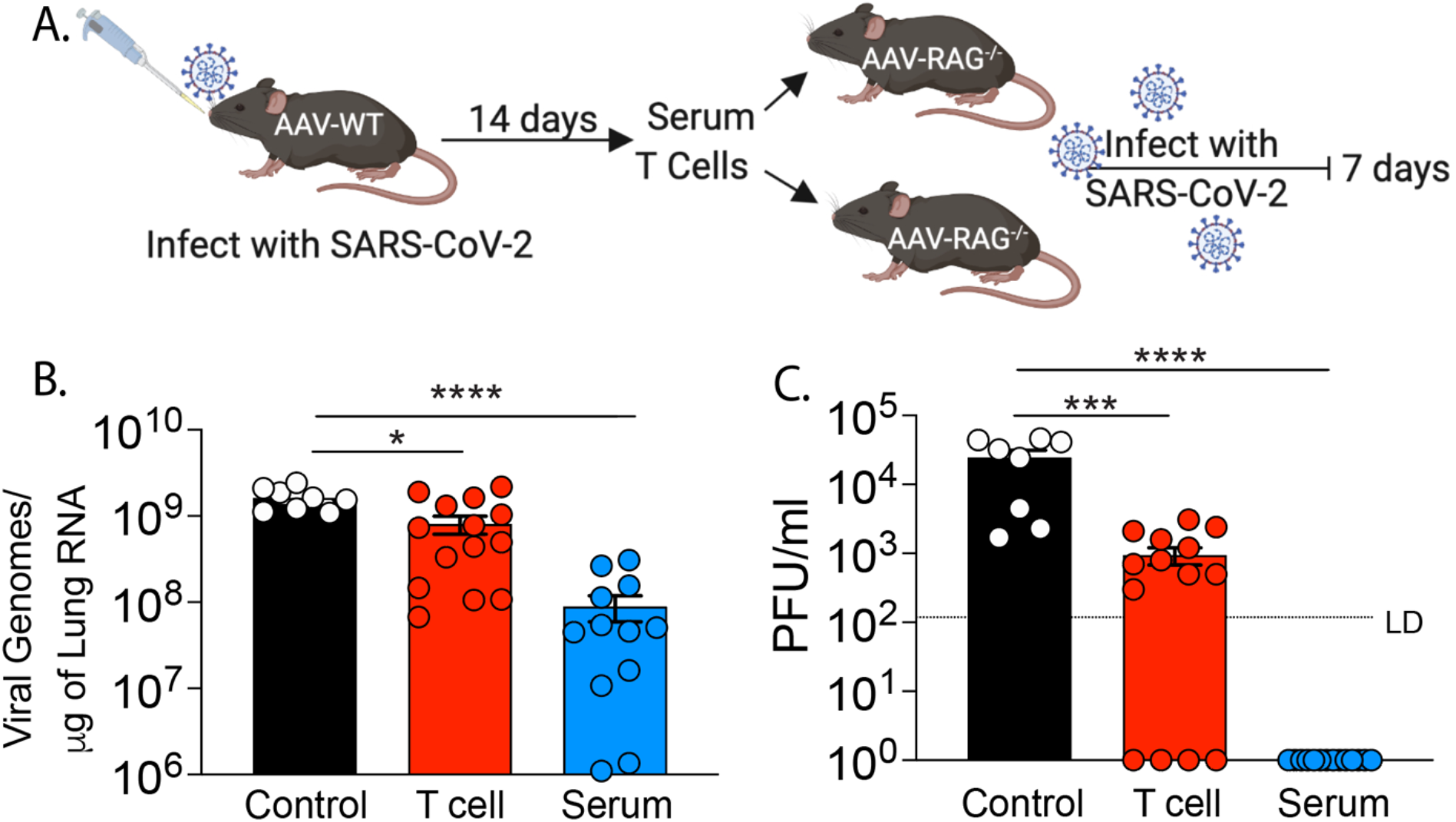
Infection of RAG^-/-^ mice after adoptive transfer of serum or T cells. (A) Experimental design. AAV-hACE2 transduced WT mice were infected with 10^6^ PFU WA1 strain SARS-CoV-2 via intranasal route. 14 DPI serum and mediastinal lymph node were collected. Total T cells were isolated by magnetic separation. AAV-hACE2 transduced RAG^-/-^ mice received adoptive transfer of PBS (control), serum (200ml), or 2×10^6^ total T cells at one day prior to infection with 10^6^ PFU WA1 strain SARS-CoV-2. Lungs were collected at 7 DPI and assessed for (B) viral RNA by quantitative PCR and for (C) viral titer by plaque assay. Individual values noted as dots and bars as mean ± SEM from 8-12 samples of two independent experiments. P values were calculated by two-tailed Mann-Whitney test. *, P< 0.05; **, P < 0.01; ***, P < 0.001, ****, P < 0.001

Finally, we assessed the role of humoral and cellular immunity in both vaccinated and convalescent mice in the protection from homologous and heterologous viral challenge. For these experiments, we utilized the K18 hACE2 mice (transgenic mice that express human ACE2) model, as they represent a lethal disease model of SARS-CoV-2 (*24, 25*). To initially characterize humoral and cellular responses to natural infection and to mRNA vaccine, mice were either infected with 500 PFU (~LD50) of SARS-CoV-2 WA1 strain once, or received 1μg mRNA Pfizer–BioNTech COVID-19 vaccine (Fig 4A) (*26*). At 14 days post immunization or infection, mice were assessed for anti-Spike S1 IgG and spike specific CD8 T cells using S539-546 epitope MHC I tetramer staining from lung (*27, 28*). We found that both groups developed similar levels of spike specific IgG (Fig 4D). Using IV labeling, we identified that both groups developed similar levels of circulating (IV^+^) spike specific CD8 T cells (CD8^+^Tetramer^+^) (Fig 4B,C). Conversely, we found that only convalescent mice developed high levels of SARS-CoV-2 specific lung resident (IV^-^) memory T cells (T_RM_) (CD8^+^CD69^+^CD103^+^Tetramer^+^), while vaccinated mice did not (Fig 4B,C). We therefore hypothesized that lung CD8^+^ T_RM_ memory cells may play a more important role in protection of convalescent mice than in vaccinated mice given their localization at the site of infection.

**Figure 4:**
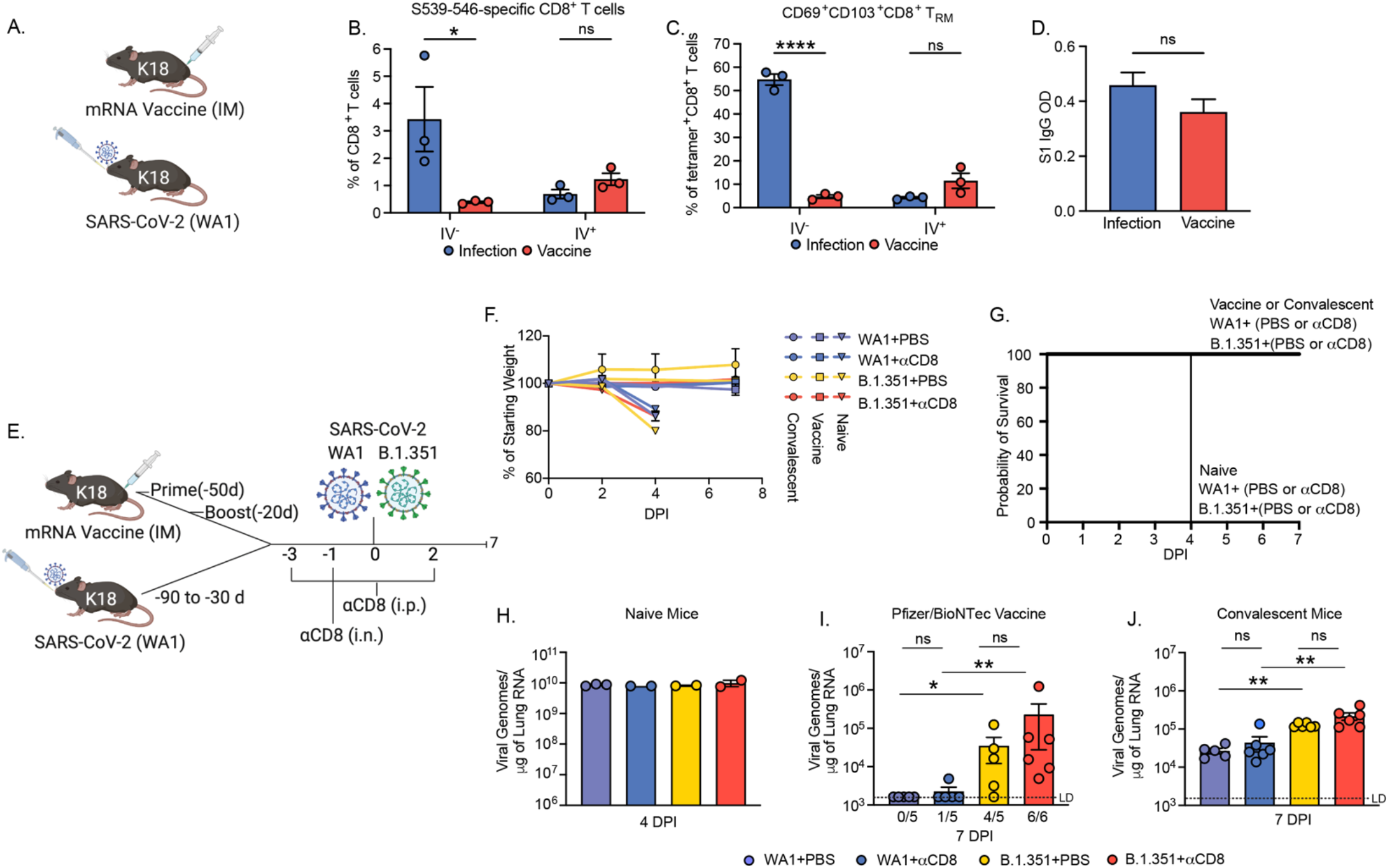
Response to homologous and B.1.351 challenge of convalescent and mRNA vaccinated mice. (A) B6.Cg-Tg(K18-ACE2)2Prlmn/J (K18-hACE2) mice received 1μg Pfizer/BioNTech mRNA vaccination via intramuscular injection or were infected with 500 PFU WA1 strain SARS-CoV-2 via intranasal administration. Two weeks post vaccination or infection, mice were injected intravenously (IV) with anti-CD45 labeling antibodies to distinguish circulating from tissue-resident T cell responses. (B) Lung-resident (IV^-^) and circulating CD8 T cells (IV^+^) specific for spike S539-546 were assessed by flow cytometry. (C) Memory T cells were assessed using CD69 and CD103 expression on lung-resident (IV^-^) and circulating (IV^+^) CD8 T cells specific for spike S539-546 by flow cytometry. (D) End dilution ELISA (1:30K) O.D. 450 IgG against SARS-CoV-2 S1. Individual values noted as dots and bars as mean ± SEM from n=3 samples. (E) Experimental design. K18-hACE2 mice either vaccinated with 1μg Pfizer/BioNTech mRNA via intramuscular injection via prime/boost with a 30 day interval, or infected with 500 PFU WA1 strain SARS-CoV-2 via intranasal administration once. Mice were injected (i.p.) with 200μg anti-mouse CD8a (clone 2.43) or PBS (control) at 3 and 1 days prior to infection, and 2 days post infection. mice received 100μg anti-mouse CD8a or PBS via intranasal administration at 1 day prior to infection. Mice were infected with 4×10^5^ PFU of with homologous strain (WA1) or VOC (B.1.351) via intranasal route. (F) Weight loss and (G) Kaplan Meier curve. Viral RNA was assessed by quantitative PCR in (H) naïve mice at 4DPI or 7DPI in (I) vaccinated or (J) convalescent mice. Values noted below X-axis (I) indicate numbers of samples tested positive/number of samples. LD (limit of detection). Individual values noted as dots and bars as mean ± SEM from n=2-6 samples. P values were calculated by two-tailed Mann-Whitney test. *, P< 0.05; **, P < 0.01; ***, P < 0.001, ****, P < 0.001.

To assess this hypothesis and further understand the role of cellular and humoral memory in protection generated by both vaccination and natural infection, we performed variant viral challenge experiments with vaccinated and convalescent mice in the setting of local and systemic CD8^+^ T cell depletion. In these experiments, the vaccinated group received 1μg mRNA Pfizer–BioNTech COVID-19 vaccine twice separated by 30 days via intramuscular injection, and then were infected with SARS-CoV-2 ~3 weeks post boost. The convalescent group had been infected with LD50 (~500 PFU) of WA1 strain 1-3 months prior to reinfection. Mice from the convalescent and vaccinated groups were then treated with CD8-depleting antibody or with PBS via systemic (intraperitoneal) administration at −3, −1, and +2 days relative to infection and via local (intranasal) administration −1 day prior to infection (Fig 4E). Mice were infected with 4×10^5^ PFU of either homologous strain (WA1) or a variant of concern (B.1.351) known to evade antibody neutralization from both convalescent and vaccinated people (*16–19*). We found that all the mice were resistant to disease, showing no physical signs of disease or weight loss (Fig 4F,G). By contrast, naïve mice infected with either WA1 or B.1.351 developed significant weight loss and visible signs of sickness, including reduced activity and responsiveness, succumbing to infection by 4 DPI (Fig 4F,G). Additionally, these two viruses seemed to replicate with similar efficacy in naïve mice as indicated by equal RNA levels at 4 DPI (Fig 4H). At 7 DPI in vaccinated animals, homologous WA1 challenge resulted in 0/5 PBS treated and 1/5 aCD8 treated animals having detectable lung viral RNA, while challenge with B.1.351 resulted in 4/5 PBS treated and 6/6 αCD8 treated animals with detectable lung viral RNA (Fig 4I). Reinfection of convalescent mice resulted in low levels of detectable viral RNA in all samples. There was increased viral RNA in the B.1.351 infected mice relative to WA1,4.7x in PBS treated and 5.0x αCD8 treated (Fig 4J). Of note, either vaccination or previous infection resulted in protection from diseases and significant reduction in viral replication by either homologous or B.1.351 challenge. These data show that mice infected with B.1.351 either in the vaccinated or the convalescent group had significantly higher levels of viral RNA than mice infected with WA1. Additionally, the mice treated with αCD8 sustained similar levels of viral RNA compared to PBS treated animals infected by the same virus. We found this to be the case in either vaccinated or convalescent mice, suggesting that memory T cells, either circulating or resident within the lung, do not play a significant role in protection conferred by natural infection or vaccine-induced immunity. These results are consistent with our adoptive transfer experiments (Fig 3) where we only observed a mild reduction of SARS-CoV-2 in T cell treated animals, while serum treatment led to significant reduction. Thus, our results indicate that both vaccination and prior infection provide significant protection against infection and complete protection against disease by B.1.351 VOC, independently of CD8 T cells.

Here we describe our use of mouse models of SARS-CoV-2 infection to identify components of adaptive immunity responsible for both viral clearance during primary infection and protection from reinfection. We found that, similar to COVID-19 patients, mice require an adaptive immune response to clear SARS-CoV-2. This indicates that in this model, innate immunity is insufficient to clear infection, which may be due to antagonism of interferon pathway by the virus (*29*). We found that either cellular or humoral arms of the adaptive immune system is sufficient to promote viral clearance, as either B cell deficient mice or mice treated with αCD8 or αCD4 antibodies alone can clear SARS-CoV-2 infection, albeit slower than in the setting of a fully competent adaptive immune system. However, depletion of both CD4 and CD8 T cells in WT mice or depletion of CD8 T cells in μMT mice led to the inability to clear the virus during primary infection. We also found that the major contribution of CD4 T cells to viral clearance during acute infection is likely the promotion of antibody production, as treatment of μMT mice, devoid of antibodies, with αCD4 only led to a small increase in viral RNA. These results indicate that T_H_1 and CD4^+^-cytotoxic T cell functions do not play a major role in viral clearance during primary infection. Consistent with these results, antigen specific CD4^+^ T cell profiling of acute and convalescent COVID-19 patients has shown that circulating T follicular helper cells dominate CD4^+^ T cell subsets, indicating the importance of CD4^+^ T cells in antibody production and clearance of acute infection. COVID-19 patient studies have shown that antigen specific CD4^+^ T cells can be detected as early as 2-4 days from symptom onset, and their early detection was found to be associated with improved outcomes and viral clearance (*30, 31*). These data clearly show that both humoral and cellular adaptive immunity contribute to clearance of SARS-CoV-2 during primary infection, and are consistent with recent patient studies showing a correlation between clinical outcome and a robust coordinated adaptive response requiring CD4^+^ T cells, CD8^+^ T cells, and antibodies (*30, 32*).

Adaptive immune memory has been detected for as long as 8 months after primary infection with SARS-CoV-2 consisting of memory CD4^+^ T cells, CD8^+^ T cells, B cells and antibodies (*1, 4, 33, 34*). Epidemiologic evidence also indicates that immune memory is sufficient to protect against reinfection in patients who have seroconverted, and mRNA-based vaccines have also show >90% efficacy in preventing COVID-19 in both phase III clinical trials and in real world settings (*3, 13–15*). As both vaccines and natural infection induce multiple types of immune memory and because reinfections in humans have been so rare, identifying the distinct components of adaptive immune memory which confer protection remain unknown.

While protective immunity after primary infection or vaccine has been shown in multiple animal models, few have identified correlates of protection (*5–12*). Many of these studies have also shown that adoptive transfer of serum from convalescent or vaccinated animals or from humans confers protection. However, the role that T cells play in conferring protection either from vaccination or natural infection has received less attention than antibodies until recently. To begin to assess the individual components of adaptive immunity that confer protection, McMahan et. al.(*6*) recently showed that macaques treated with CD8-depleting antibody had higher RNA viral loads in nasal swab at 1 DPI than mock treated animals. However, this difference normalized by 2 DPI, and no difference at any time post infection was noted in BAL, suggesting that CD8 T_RM_ may only confer added protection in the upper respiratory tract of macaques. In mouse models, nucleocapsid (N) based vaccines, which likely rely mostly on T cell-based protection, have shown differing results. Matchett et. al.(*35*) showed that IV administration of Adenovirus 5 vector expressing N conferred protection, while Dinnon et. al. (*36*) showed no protection via foot pad injection of Venezuelan equine encephalitis virus replicon particle expressing N.

To expand upon these studies in identifying the individual roles of humoral and cellular immunity in protection, we preformed adoptive transfer studies in mice deficient in adaptive immunity (RAG1^-/-^) and found that both T cells and convalescent serum from previously infected animals could reduce viral load. However, only serum was able to achieve viral clearance. Consistent with McMahan et. al., we showed that CD8 memory T cells, either T_RM_ or circulating memory cells, are not required to confer significant protective immunity in the lower respiratory tract in either convalescent or vaccinated animals. It is certainly possible that their effect may be obscured by an overwhelming humoral response; however, even in the setting of decreased humoral protection (challenge with VOC B.1.351) we did not see a significant loss of protection in CD8^+^ T cell depleted animals indicated by either disease or viral load. These data support antibody responses as the key determinant of protection against SARS-CoV-2 reinfection, with CD8^+^ memory T cells playing only a supporting role.

SARS-CoV-2 variants that significantly evade humoral immunity have recently been identified. The variant which has consistently shown the strongest ability to evade humoral responses in *in vitro* neutralization assays is B.1.351. We found that while there was some loss of protection against infection in either vaccinated or convalescent mice, humoral immunity was sufficient to completely protect from disease in mice infected by B.1.351. These data build upon a recent case control study that reported increased rates of vaccine breakthrough with B.1.351 in fully vaccinated subjects, though most cases were asymptomatic (*16*). However, the degree to which mRNA vaccinated people are susceptible to clinical disease by B.1.351 remains unclear.

In conclusion, we provide insights into both the immunologic determinants of viral clearance and protection. While T cells were important in the clearance of primary infection, they were not required for protection against reinfection or vaccine-mediated protection. These results are reassuring as they indicate that a robust humoral immune response is sufficient even in the setting of decreased neutralizing capacity. These results also have important public health and vaccine development implications, as they suggest that antibody mediated immunity may be a sufficient correlate of protect.

## Acknowledgement

We thank H. Dong and M. Linehan for technical and logistical assistance. We thank Patrick Roberts and Bryan Cretella from the Yale Health Pharmacy for providing residual vaccine used in this study. We thank Craig Wilen for technical assistance and providing reagents. We thank Carolina Lucas for technical expertise and for critical feedback on the manuscript. We thank Barney Graham (NIH-VRC) for kindly providing VeroE6 cells overexpressing ACE2 and TMPRSS2. We thank the NIH Tetramer Core Facility for providing PE labelled SARS-CoV-2 S 539-546 tetramer (H-2K(b)). We also give special recognition of the services of Ben Fontes and the Yale EH&S Department for their on-going assistance in safely conducting biosafety level 3 research.

## Funding

Fast Grant from Emergent Ventures (AI)

Mercatus Center (AI)

National Institutes of Health grant R01AI157488 (AI)

National Institutes of Health grant T32AI007517 (BI)

Howard Hughes Medical Institute (AI)

## Author contributions

Conceptualization: BI, SBO, and AI

Methodology: BI and AI

Investigation: BI, TM, JK, ES, and BM

Visualization: BI and TM

Funding Acquisition: AI

Writing- original draft: BI and AI

Writing- review & editing: BI, TM, JK, ES, SBO, AI

## Data and materials availability

All data contained in this paper are publicly available upon request.

